# Genome-wide analysis of long noncoding RNAs, 24-nt siRNAs, DNA methylation and H3K27me3 marks in *Brassica rapa* and related species

**DOI:** 10.1101/2020.11.06.370940

**Authors:** Hasan Mehraj, Daniel J. Shea, Satoshi Takahashi, Naomi Miyaji, Ayasha Akter, Motoaki Seki, Elizabeth S. Dennis, Fujimoto Ryo, Kenji Osabe

**Affiliations:** Graduate School of Agricultural Science, Kobe University, Rokkodai, Nada-ku, Kobe 657-8501, Japan; Bloomberg LP, Chiyoda-ku, Tokyo 100-0005, Japan; Riken center for Sustainable Resource Science, Yokohama, Kanagawa, 230-0045, Japan; RIKEN Cluster for Pioneering Research, 2-1 Hirosawa, Wako, Saitama, 351-0198, Japan; Kihara Institute for Biological Research, Yokohama City University, 641-12 Maioka-cho, Totsuka-ku, Yokohama, Kanagawa 244-0813, Japan; CSIRO Agriculture and Food, Canberra, ACT 2601, Australia; University of Technology, Sydney, PO Box 123, Broadway, NSW 2007, Australia; Institute of Scientific and Industrial Research, Osaka University, Mihogaoka, Ibaraki-city, Osaka, Japan

**Keywords:** Brassica rapa, noncoding RNA, siRNA, DNA methylation, H3K27 methylation, epigenome, Brassica nigra, Brassica oleracea, Brassica juncea, Brassica napus

## Abstract

Long noncoding RNAs (lncRNAs) are RNA fragments that generally do not code for a protein but are involved in epigenetic gene regulation. In this study, lncRNAs of *Brassica rapa* were classified into long intergenic noncoding RNAs, natural antisense RNAs, and intronic noncoding RNAs and their expression were analyzed in relation to genome-wide 24-nt small interfering RNAs (siRNAs), DNA methylation, and histone H3 lysine 27 trimethylation marks (H3K27me3). More than 65% of the lncRNAs analyzed consisted of one exon, and more than 55% overlapped with inverted repeat regions (IRRs). Overlap of lncRNAs with IRRs or genomic regions encoding for 24-nt siRNAs resulted in increased DNA methylation levels and when both were present, there were further increase in DNA methylation levels. LncRNA did not overlap greatly with H3K27me3 marks, but the expression level of intronic noncoding RNAs that did coincide with H3K27me3 marks was higher than without H3K27me3 marks. The Brassica genus comprises important vegetables and oil seed crops grown across the world. *Brassica rapa* is a diploid (AA genome) thought to be one of the ancestral species of both *B. juncea* (AABB genome) and *B. napus* (AACC) through genome merging (allotetrapolyploidization). Complex genome restructuring and epigenetic alterations are thought to be involved in these allotetrapolyploidization events, but the detailed mechanism is not known. Comparison of lncRNAs between *B. rapa* and *B. nigra, B. oleracea, B. juncea*, and *B. napus* showed the highest conservation with *B. oleracea*. This study presents a comprehensive analysis of the epigenome structure of the *B. rapa* at multi-epigenetic levels (siRNAs, DNA methylation, H3K27me3, and lncRNAs) and offers insights into the function of lncRNA in the Brassica genus.

## Introduction

The Brassica genus comprises vegetable and oil seed crops. The “Triangle of U” proposed the genomic relationship among six major species of the Brassica genus. Three allotetraploid species, each of which contains two complete diploid genomes derived from two different parental species, *Brassica juncea* L. (AABB genome, 2n = 4x = 36), *B. napus* L. (AACC, 2n = 4x = 38), and *B. carinata* L. (BBCC, 2n = 4x = 34) are derived from the natural hybridization of the diploid species, *B. rapa* L. (AA, 2n = 2x = 20), *B. nigra* L. (BB, 2n = 2x = 16), and *B. oleracea* L. (CC, 2n = 2x = 18) [1]. Some species in the Brassica genus show morphological divergence (termed morphotype). *B. rapa* includes leafy vegetables including Chinese cabbage (var. *pekinensis*), pak choi (var. *chinensis*), and komatsuna (var. *perviridis*), root vegetables including turnip (var. *rapa*), and oilseed crops (var. *oleifera*) [2]. The first whole genome sequence determined in the genus Brassica was that of *B. rapa* [3]. Later the whole genome sequences of *B. oleracea, B. nigra, B. napus*, and *B. juncea* were determined [4–7].

Plant transcriptome analyses have revealed transcribed RNAs devoid of protein-coding potential, which are called noncoding RNAs (ncRNAs). Some ncRNAs contain exons, which potentially code for short proteins or peptides, however, experimental investigation is required to validate their functions. Two families of ncRNAs are known; long noncoding RNAs (lncRNAs) that are longer than 200 nucleotides (nt), and small RNAs (sRNAs) that are ∼18-30 nt in length [8–12]. LncRNAs are classified by their position and orientation of transcription; long intergenic noncoding RNAs (lincRNAs), intronic noncoding RNAs (incRNAs) derived from introns, and natural antisense transcripts (NATs) transcribed from the complementary DNA strand of their associated genes [8-12]. Recent studies have shown that lncRNAs play crucial roles in various physiological processes such as vernalization [13-15], photoperiod-sensitive male sterility [16], red-light-mediated seedling photomorphogenesis [17], seed dormancy [18], and the transcriptional regulation of plant innate immunity [19].

Epigenetics has been defined as ‘‘the study of changes in genome expression that are mitotically and/or meiotically heritable and that do not entail a change in DNA sequence’’ [20]. DNA methylation and histone modification are well-known epigenetic modifications, and lncRNAs are considered to be involved in epigenetic regulation [21]. In plants, DNA methylation is established through RNA-directed DNA methylation (RdDM) [22,23]. Plant-specific RNA POLYMERASE IV (Pol IV) and Pol V are involved in RdDM. Pol IV-transcribed ncRNAs are cleaved into 24-nucleotide small interfering RNAs (24-nt siRNAs) by DICER-LIKE 3 (DCL3), and 24-nt siRNAs are loaded into ARGONAUTE 4 (AGO4). PolV-transcribed lncRNAs act as scaffold molecules of AGO-siRNA complexes, which recruit DOMAINS REARRANGED METHYLTRANSFERASE 2 (DRM2) for catalyzing *de novo* DNA methylation. In *Arabidopsis thaliana* two cold-induced lncRNAs (COLDAIR and COLDWRAP) regulate histone methylation by recruiting polycomb repressive complex 2 (PRC2), which catalyzes the tri-methylation of histone H3 lysine 27 (H3K27me3), to the chromatin region of *FLOWERING LOCUS C* (*FLC*) [14,24-26].

Previous studies have identified lncRNAs associated with different lines, tissues or environmental changes in Brassica [27–33]. However, there are few reports of the association between lncRNAs and epigenetic modifications in *B. rapa*. In this study, we examined the DNA methylation levels and H3K27me3 levels in the region covering lncRNAs, and found an association between lncRNAs and inverted repeat regions (IRRs) or 24-nt siRNAs, but association with H3K27me3 to a specific type of lncRNA.

## Materials and methods

### Plant materials and growth conditions

Six lines of *B. rapa*, inbred lines of RJKB-T24 (var. *pekinensis*) [34] and Yellow sarson (var. *trilocularis*), doubled haploid lines of BRA2209 (var. *rapa*), Homei (var. *pekinensis*), and Osome (var. *perviridis*), and a commercial F_1_ hybrid cultivar, ‘Harunosaiten’ (var. *pekinensis*) (Watanabe Seed Co., Ltd.) were used. Three *B. oleracea* cabbage (var. *capitata*) F_1_ hybrid cultivars, ‘Reiho’, ‘Matsunami’ (Ishii Seed Growers CO., LTD), and ‘Kinkei 201’ (Sakata Seed Co., Ltd.) were also used.

Seeds were surface sterilized and grown on agar solidified Murashige and Skoog (MS) plates with 1 % (w/v) sucrose under long day (LD) condition (16h light / 8h dark) at 21 °C. Fourteen-day first and second leaves of *B. rapa* and 19-day first and second leaves of *B. oleracea* were harvested for isolation of genomic DNA or total RNA.

RJKB-T24 was used for RNA-sequencing (RNA-seq) for detection of lncRNAs. Briefly, after performing QC of the sequenced reads, putative mRNAs were identified by aligning the sequence reads to the *B. rapa* reference genome v1.5 using HISAT2 and then assembling transcripts with Stringtie. Assembled transcripts with a mapping code of ‘u’, indicating they are intergenic but not part of the annotated reference genome, were then compared to the SwissProt database using blastx (e-value 1e-10). Transcripts with hits were classified as putative mRNAs, while transcripts with no hits were classified as putative lincRNAs. A more detailed description of this method can be found in Shea et. al, 2018 [27].

### DNA extraction and PCR

Genomic DNA was isolated by the Cetyl trimethyl ammonium bromide (CTAB) method [35]. The PCR reaction was performed using the following conditions; 1 cycle of 94 °C for 3 min, 35 cycles of 94 °C for 30 s, 55 °C for 30 s, and 72 °C for 1 min, and final extension at 72 °C for 3 min. The PCR products were electrophoresed on 1.0% agarose gel. Primer sequences used in this study are shown in S1 Table.

### RNA extraction and RT-PCR

Total RNA from the first and second leaves were isolated by SV Total RNA Isolation System (Promega Co., WI, USA). To analyze lncRNA expression, cDNA was synthesized from 500 ng total RNA using PrimeScript RT reagent Kit (Takara Bio., Shiga, JAPAN). The absence of genomic DNA contamination was confirmed by PCR using a control without reverse transcriptase. The PCR conditions were 94 °C for 2 min followed by 35 cycles of 94 °C for 30 s, 55 °C for 30 s, and 68 °C for 30 s. The primers used in this study are listed in S1 Table.

### Detection of epigenetic states in lncRNA regions

To examine the epigenetic states (DNA methylation levels, H3K27me3, and 24-nt siRNA levels) of lncRNA encoding regions in *B. rapa*, we used previous sequence data of whole genome bisulfite sequencing (WGBS) [36], chromatin immunoprecipitation sequencing (ChIP-seq) [37], and small RNA-sequencing (sRNA-seq) data [36], which were generated using samples from the same line, tissue, and developmental stages but harvested independently.

The reads of WGBS were mapped to the *B. rapa* reference genome v.1.5 using Bowtie2 version 2.2.3 and Bismark v0.14.3, and data covering genomic regions encoding lncRNA in chromosomes A01 to A10 were extracted. In order to estimate the methylation levels of CG, CHG, and CHH contexts, the numbers of methylated and unmethylated reads were extracted for each cytosine position using bismark methylation extractor script with the paired-end parameter. The methylation level at each cytosine position was calculated by dividing the number of methylated cytosines (mC) reads by the total number of reads.

The reads of sRNA-seq were mapped to the *B. rapa* reference genome v.1.5 using Bowtie2 version 2.2.3. We classified the alignment reads by length, and the 24-nt aligned reads covering the lncRNA regions in chromosomes A01 to A10 were extracted.

The reads of ChIP-seq using anti-H3K27me3 (Millipore, 07-449) antibodies were mapped to the *B. rapa* reference genome v.1.5 using Bowtie2 version 2.2.3, and data covering genomic regions encoding for lncRNA in chromosomes A01 to A10 were extracted.

### Comparison of putative mRNA and lncRNAs in *B. rapa* to other related species of *Brassica*

The putative mRNAs and lncRNAs were first compared to the reference genomes of *B. nigra, B. oleracea, B. juncea*, and *B. napus* by best-hit blastn (e-value 1e-10) to identify homologous regions in closely related Brassica species [6,7,38]. In order to parse the local High-scoring segment pair (HSP) alignments produced by blastn, genBlastA was used to produce a representative putative gene that is homologous to the query [39]. The parsed local alignments were then analyzed by a custom python script (available at http://www.github.com/danshea/lncRNA) to examine the overall alignment length and computed coverage of the aligned homologous region to the putative mRNA or lncRNA query transcript sequence. These results were then imported into R and plotted to assess overall relative coverage of the sequences among the Brassica species.

## Results

### Characterization of lncRNA in *B. rapa*

We identified 1,444 lincRNAs, 551 NATs, and 93 incRNAs using RNA-seq data of 14-day first and second leaves with and without four weeks of cold treatment [27]. In order to investigate the relationship between lncRNAs and epigenetic modifications or the species specificity of lncRNAs, we analyzed in more detail the RNA-seq data of the 14-day first and second leaves without cold treatment. There was no strong bias in the chromosomal distribution of expressed lncRNAs (S1 Fig). More than 65% of lncRNAs contained one exon (lincRNAs, 65.7%; NATs, 72.4%; incRNAs, 71.0%), whereas the proportion of mRNAs containing only one exon is 15.1% (Fig 1A). The mean transcript lengths of lincRNAs (725 nt) and incRNAs (779 nt) were shorter than that of NATs (1,271 nt) and mRNAs (1,305 nt, Fig 1B). About 40% of lincRNAs were located within 2 kb of the genic regions, and about 10% of lincRNAs were located more than 20kb from the genic region (Fig 1C). In this study, we focused on lncRNAs mapped to the chromosomes A01 to A10 as previous WGBS, ChIP-seq, and sRNA-seq have omitted the placed scaffolds for their analyses [36,37]. 763 of 1,173 (65.0%) lincRNAs, 291 of 529 (55.0%) NATs, and 66 of 92 (71.7%) incRNAs overlapped with IRRs such as transposable elements (TEs) detected by RepeatMasker, suggesting that IRRs are the source of more than half of lncRNAs in *B. rapa* (Fig 2).

**Fig 1.**
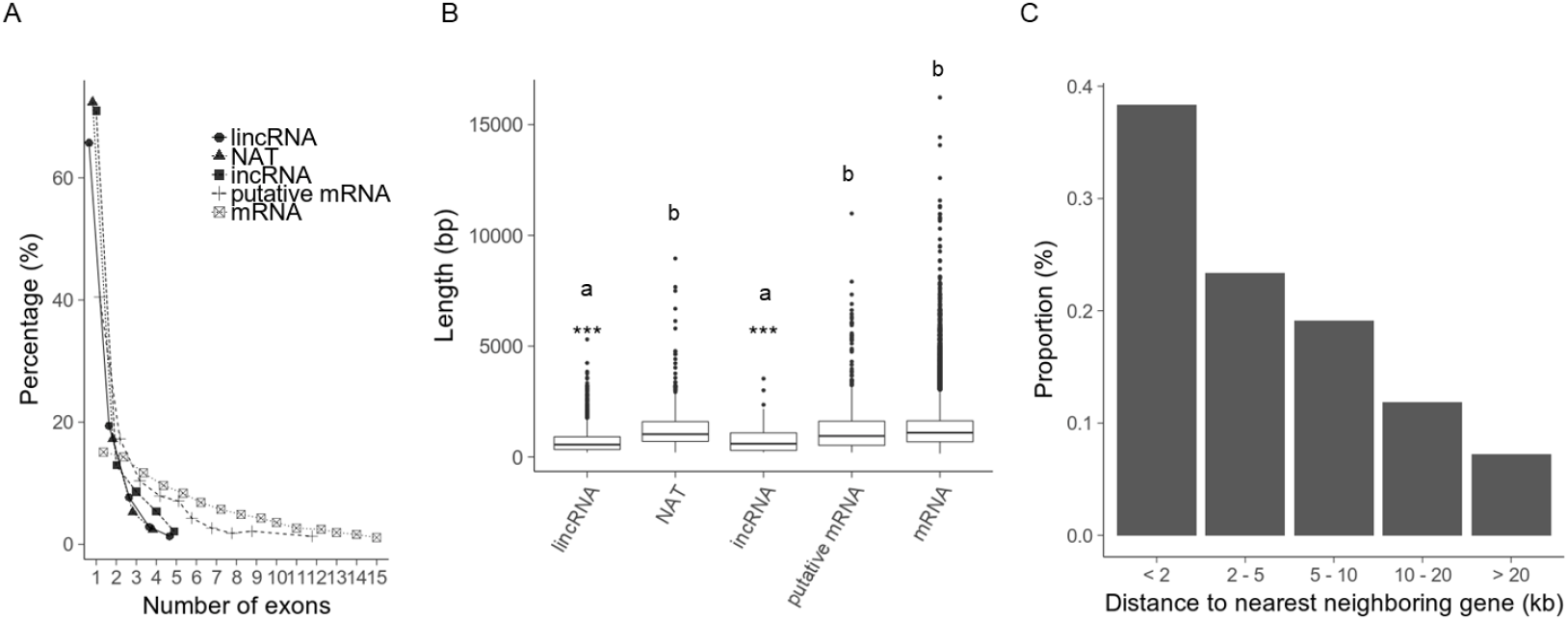
Analysis of lncRNAs of 14-day first and second leaves of *B. rapa*. (A) Number of exons in lncRNA. (B) Nucleotide length of lincRNA, NAT, incRNA, putative mRNA and mRNA. “a” and “b” represent significant differences by one-way ANOVA test (“*”, *p*<0.05; “***”, *p*<0.001). (C) The proportion lncRNA distances to the nearest gene.

**Fig 2.**
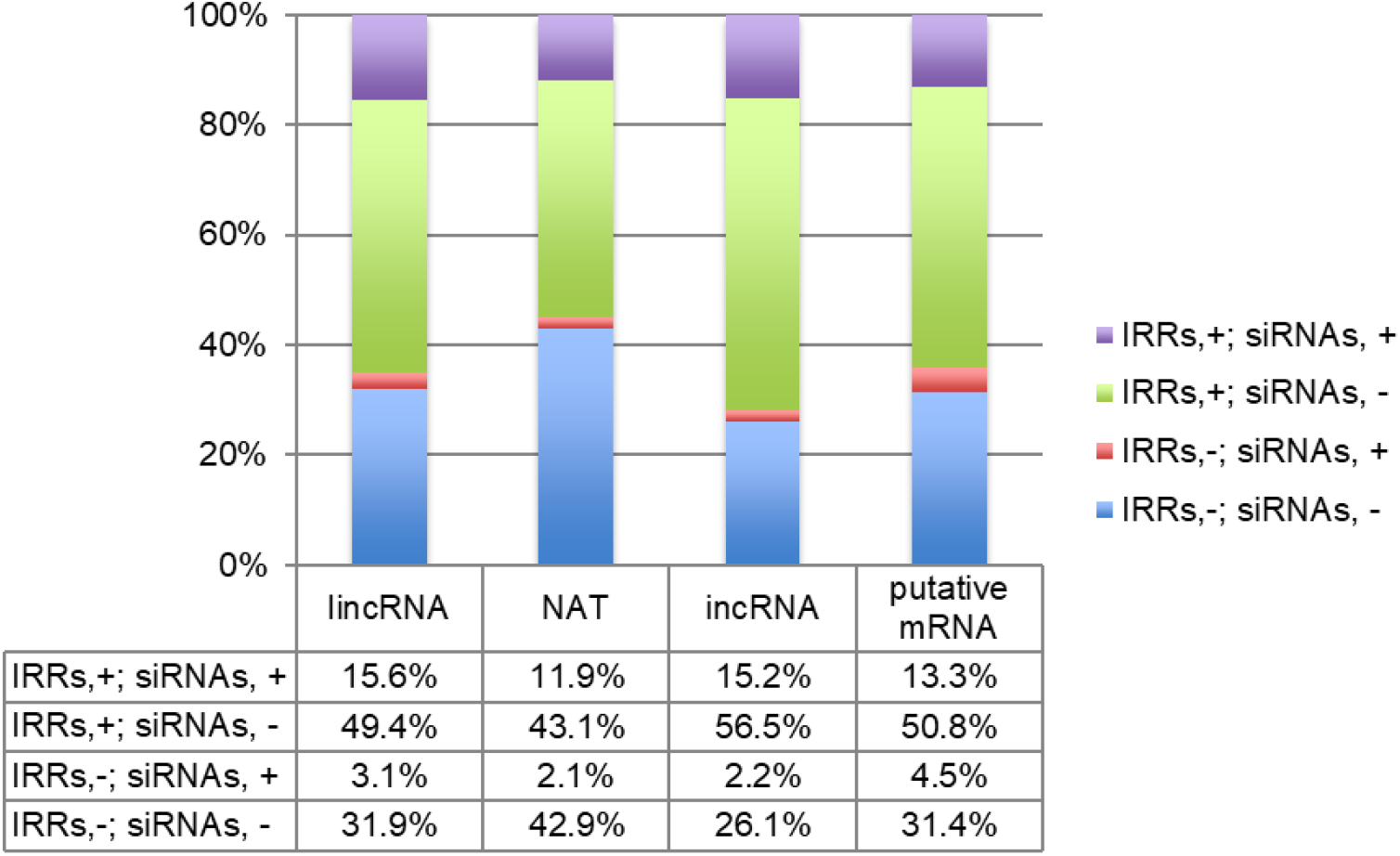
Proportion of each type of lncRNA and putative mRNAs that overlap with IRRs and siRNA. “+” indicates with and “-” indicated without overlapping IRRs or genomic regions encoding siRNAs.

### Relationship between lncRNAs and siRNA or DNA methylation

We have performed sRNA-seq using 14-day first and second leaves, which are identical developmental stages and tissues to previously analyzed samples [36], but independently harvested for this study. We identified the lncRNAs having perfect sequence identity to genomic regions encoding for 24-nt siRNAs. 219 of 1,173 (18.7 %) lincRNAs, 74 of 529 (14.0%) NATs, and 16 of 92 (17.4%) incRNAs in A01-A10 overlapped with unique-mapped genomic regions encoding for 24-nt siRNAs, and more than 80% of the lncRNAs overlapping with genomic regions encoding for 24-nt siRNAs were from regions that harbored IRRs (Fig 2). 24-nt siRNAs were mapped in a similar way to lncRNA and its 5’ and 3’ flanking regions; this mapping pattern is different from those of genic regions or IRRs (Fig 3).

**Fig 3.**
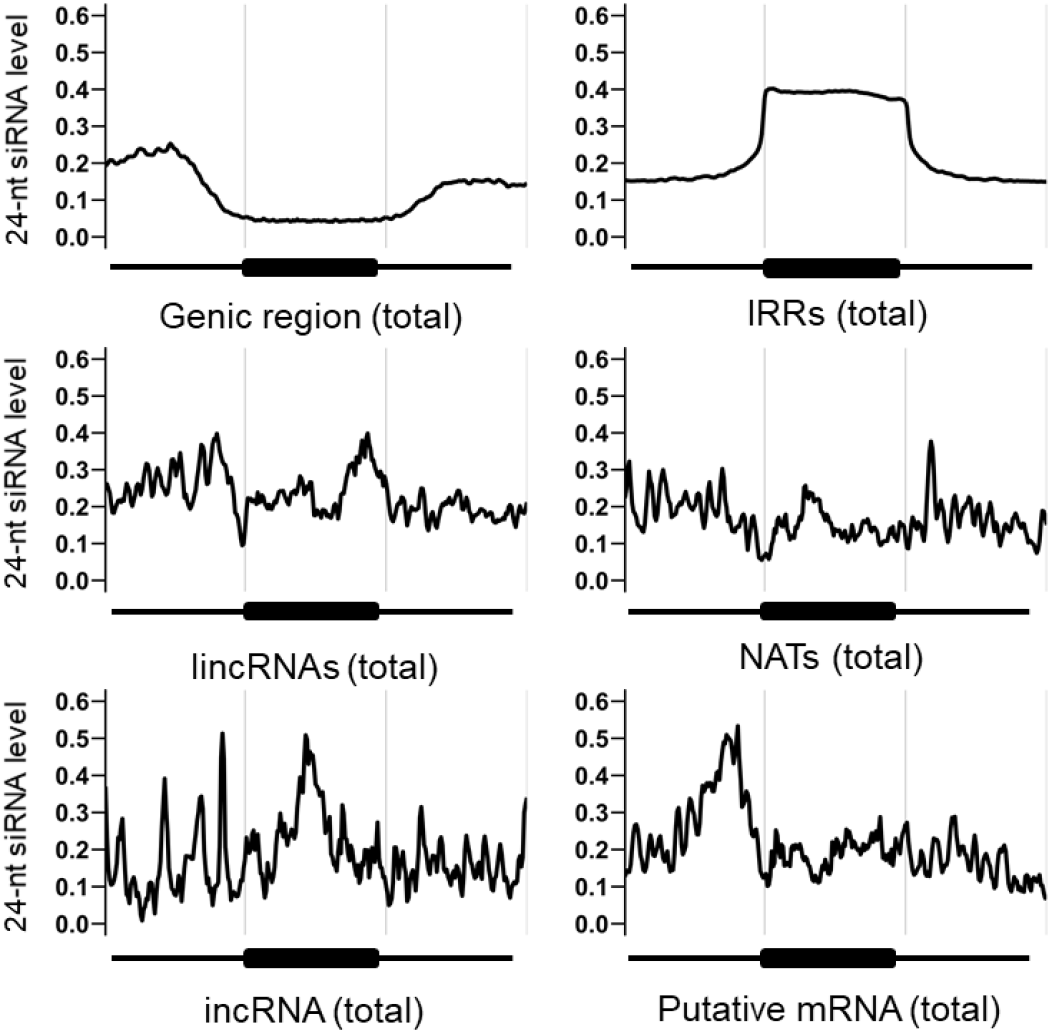
24nt-siRNAs mapped to genic region, IRRs, putative mRNAs and each type of lncRNAs. The x-axis represents the target region (genic region, IRRs, putative mRNAs, or lncRNAs) and the flanking 5’ and 3’ regions. The y-axis represents the reads per million (RPM) value of 24-nt siRNA that overlaps with the corresponding target region.

We examined the whole genome DNA methylation state by WGBS using the same 14-day first and second leaves [36]. The average DNA methylation levels in regions covering lncRNAs were similar to those of the whole genome (S2 Fig). DNA methylation level in regions producing NATs was lower than those producing any of the three types of lncRNAs. Overlap of lncRNAs with IRRs or genomic regions encoding for 24-nt siRNAs was associated with increased DNA methylation levels and overlapping with both caused further increases in DNA methylation levels (Fig 4).

**Fig 4.**
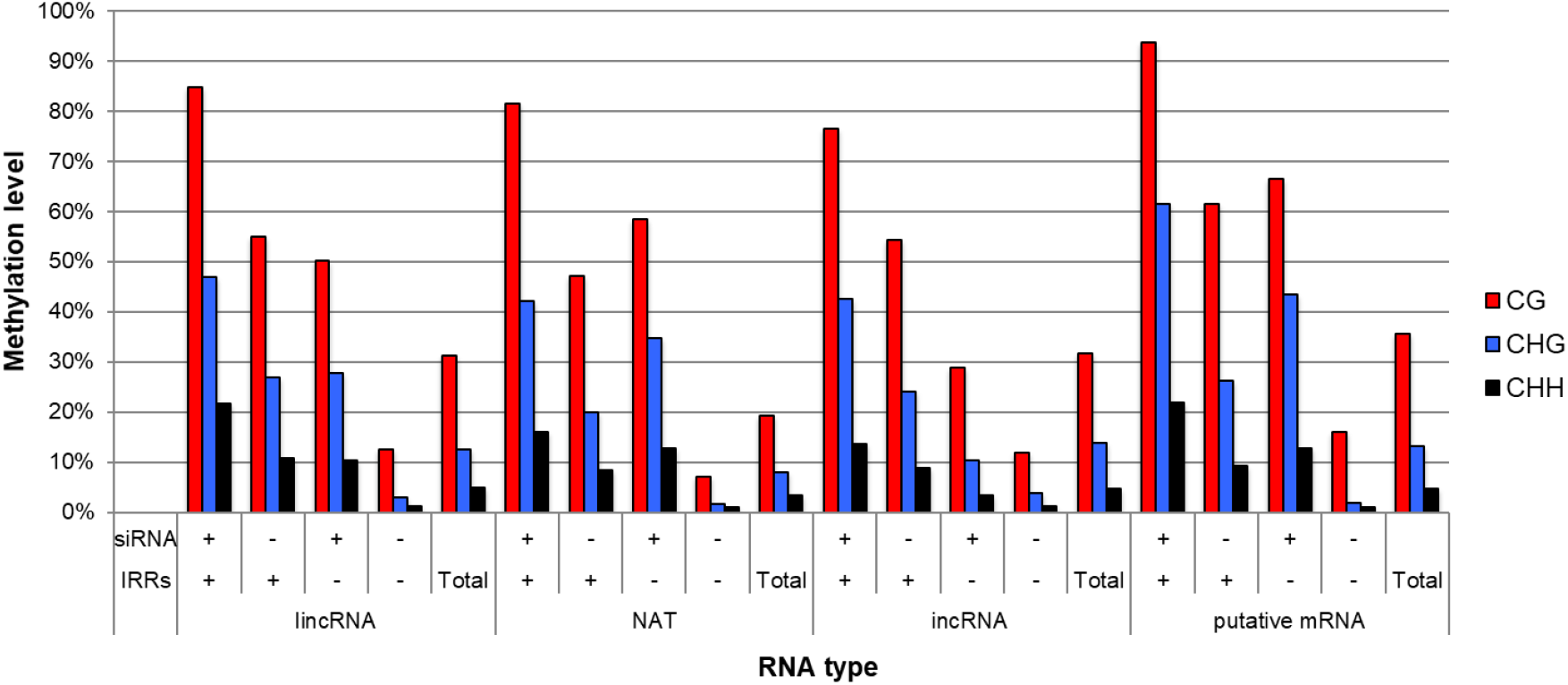
DNA methylation level of genomic regions encoding each RNA type with siRNAs or IRRs. “+” indicates with and “-” indicates without overlapping genomic regions encoding for siRNAs or IRRs.

The DNA methylation were similar in lincRNA and NAT encoding regions, and DNA methylation levels in the 5’ and 3’ flanking regions were higher than in the body regions encoding lincRNAs and NATs (S3 Fig). The DNA methylation levels over the incRNA encoding regions and the flanking regions did not change (S3 Fig). Overall, mapping of DNA methylation levels to lncRNA encoding regions were different from those of genic regions or IRRs (S3 Fig).

We examined the influence on the lncRNA expression level when IRRs or siRNA clusters on the genome spanned the lncRNA regions. IRRs did not affect the expression levels of lncRNAs (Fig 5). The expression levels of lncRNAs covered with siRNA clusters were higher than those not covered by siRNA clusters (Fig 5)

**Fig 5.**
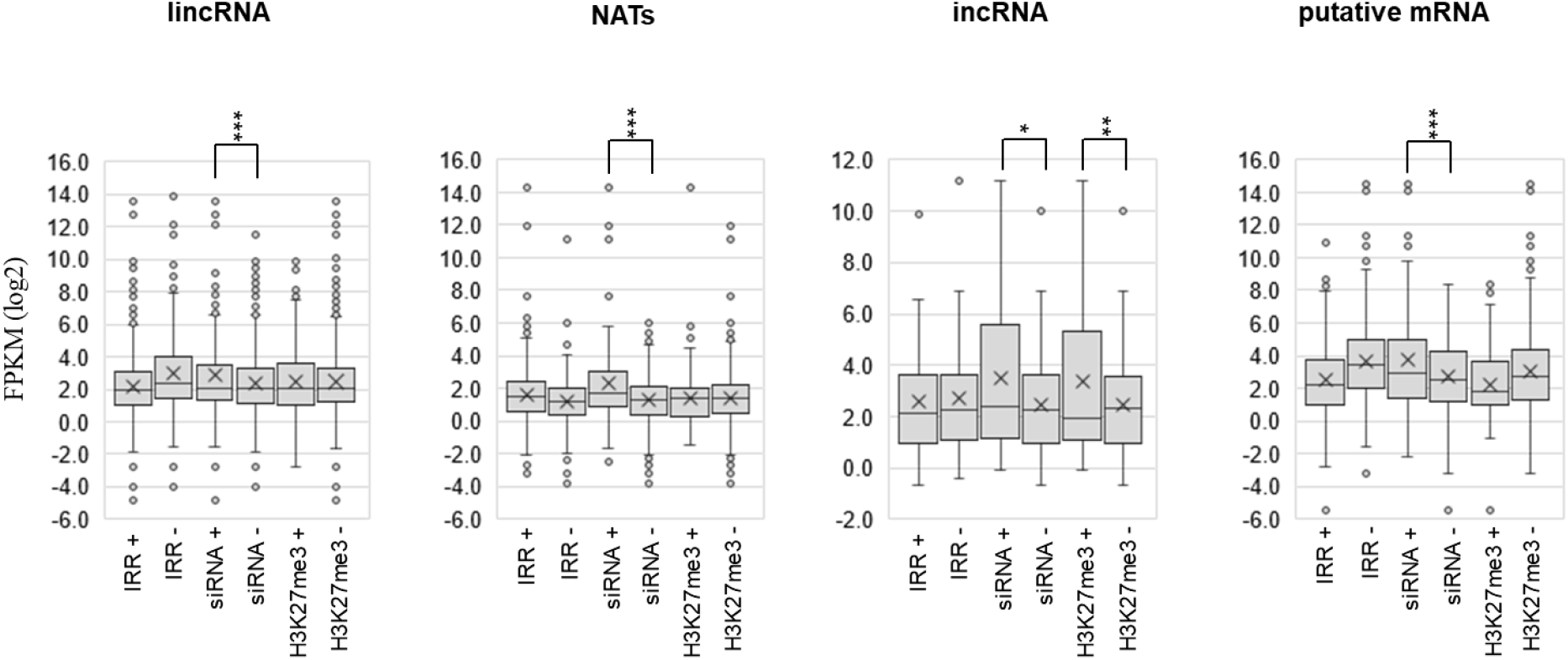
Expression level of each type of RNAs with (+) or without (-) overlapping IRRs, 24nt-siRNAs, or H3K27me3. *, *p*<0.05; **, *p*<0.01; ***, *p*<0.001 (Student t-test)

### Relationship between lncRNAs and H3K27me3

We have examined the H3K27me3 distribution using the 14-day first and second leaf samples [37] and identified the genomic regions encoding for lncRNAs that have H3K27me3 marks. 127 of 1,173 (10.8 %) lincRNAs, 83 of 529 (15.7%) NATs, and 15 of 92 (16.3%) incRNAs encoding genomic regions had H3K27me3 marks (S4 Fig). H3K27me3 was enriched in the transcribed region of lncRNAs, similar to the genic regions, but H3K27me3 levels in lncRNA encoding genomic regions were lower than in regions coding for mRNAs (S5 Fig). The expression level of incRNAs was higher when the encoding regions had H3K27me3 than without H3K27me3 (Fig 5). In NATs and lincRNAs, there was no difference of expression level with and without H3K27me3 on their encoding regions (Fig 5).

### Characterization of un-annotated genes

In 2,052 lncRNAs mapped to intergenic regions of the genome, 608 transcripts had hits (e-value < 1.0e-10) against the Swissport database using BLASTX, indicating they could be un-annotated genes. The expression levels of the putative mRNAs from these regions were similar to that of mRNA [27]. Putative mRNAs tended to have fewer exons compared with annotated mRNAs (Fig 1A). The mean transcript length of putative mRNA genes was similar to that of mRNAs (Fig 1B).

314 of 490 (64.1%) putative mRNAs in A01-A10 overlapped with IRRs (Fig 2) but the expression level of putative mRNAs overlapping with IRRs was similar to those not-overlapping with IRRs (Fig 5). Mapping of 24-nt siRNAs and DNA methylation levels to genomic regions encoding putative mRNAs were similar to that of genic regions (Fig 3, S3 Fig). The average expression level of putative mRNAs with overlapping 24-nt siRNAs was higher than that without any overlap (Fig 5). The pattern of the average of DNA methylation level in the genomic region encoding putative mRNAs and their flanking regions was similar to that to the average of the genic regions (S3 Fig). Overlapping IRRs or 24-nt siRNAs resulted in an increase in DNA methylation levels and overlapping with both causes further increase in DNA methylation levels (Fig 4).

53 of 490 (10.8%) genomic regions corresponding to putative mRNAs had H3K27me3. H3K27me3 was enriched in the genomic regions encoding for lncRNAs, especially around the transcription start site, and this was similar to the pattern in the genic region (S5 Fig). The expression levels of putative mRNAs with corresponding genomic H3K27me3 were lower than those without H3K27me3, but there was no significant difference (Fig 5).

### Examination of the conservation of lncRNAs among the Brassica genus

Using the sequences of the 2,088 lncRNAs of *B. rapa*, a best-blast hit search against the *B. nigra, B. oleracea, B. juncea*, and *B. napus* reference genomes was conducted using GenBlastA (e-value = 1e-10) to examine the conservation of lncRNAs. *B. rapa* lncRNAs were most conserved in *B. oleracea* and moderate conservation was observed in *B. nigra* and *B. napus*. LncRNAs were less conserved in *B. juncea*, especially in the incRNAs (Fig 6).

**Fig 6.**
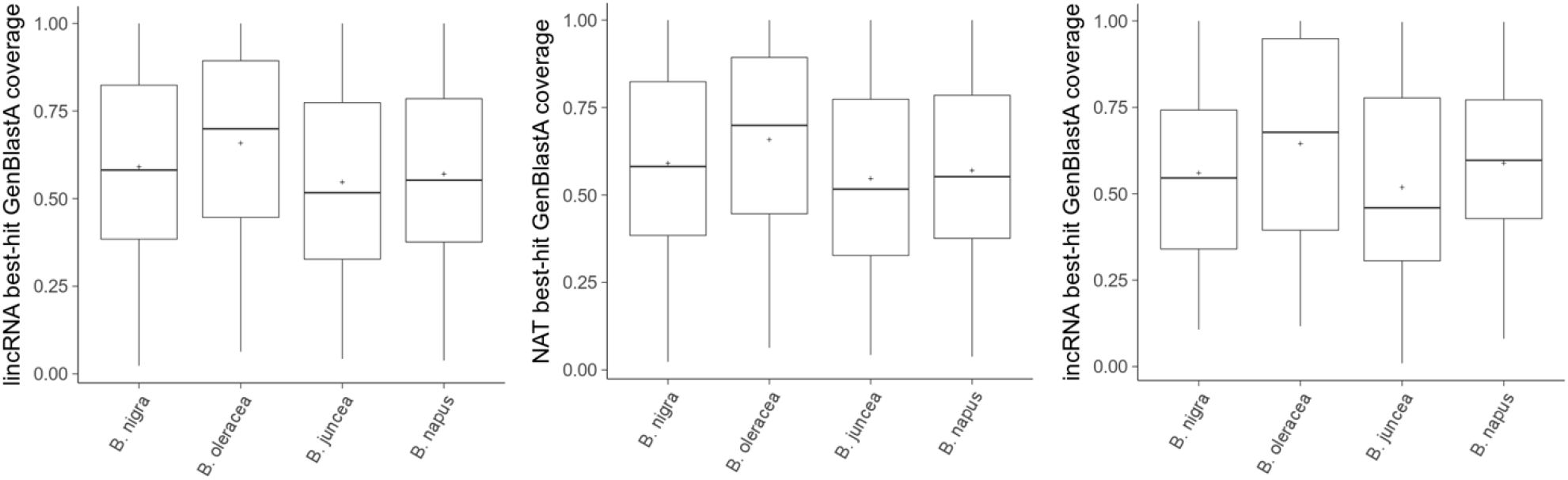
BLAST search of each type of lncRNA from *B. rapa* against *B. nigra, B. oleracea, B. juncea*, and *B. napus*.

We selected twelve lncRNAs that showed high sequence similarity with the *B. oleracea* reference genome. We tested whether these lncRNA coding genomic sequences were conserved in three commercial cultivars of cabbage (*B. oleracea*) by PCR using genomic DNA as template. PCR amplification of genomic regions corresponding to all twelve lncRNAs was confirmed in all three *B. oleracea* lines. Next, we tested by RT-PCR whether these putative lncRNAs in *B. oleracea* are expressed in the first and second leaves. Six of twelve lncRNAs were expressed in all three cultivars. One lncRNA was expressed in two of three lines, and one lncRNA was expressed in one of three lines. The remaining four lncRNAs were not expressed or were slightly expressed in all three lines (Table 1). We also examined the variation within *B. rapa* species using six lines. Seven of the twelve lncRNAs were expressed in all six lines, and the remaining five lncRNA showed line-specificity (Table 1).

**Table 1.** Conservation and diversity of the expression of lncRNAs within *B. rapa* species or between species. RT-PCR results of each lncRNAs “+++” indicates strong PCR band, “+” indicates weak PCR band, and “-” indicates no PCR band.

|  | *B. rapa* |  |  |  |  |  | *B. oleracea* |  |  |
| --- | --- | --- | --- | --- | --- | --- | --- | --- | --- |
| lncRNAs | RJKB-T23 | Homei | Harunosaiten | BRA2209 | Osome | Yellow Sarson | Reiho | Matsunami | Kinkei |
| M15784 | +++ | - | +++ | - | +++ | +++ | - | - | - |
| M26919 | +++ | +++ | +++ | +++ | +++ | +++ | +++ | +++ | +++ |
| M3316 | +++ | +++ | +++ | +++ | +++ | +++ | +++ | +++ | - |
| M491 | +++ | - | +++ | +++ | +++ | - | + | + | - |
| M17356 | +++ | + | - | - | - | - | - | - | - |
| M17153 | +++ | - | - | - | - | + | +++ | +++ | +++ |
| M25534 | +++ | +++ | +++ | +++ | +++ | +++ | +++ | +++ | +++ |
| M259 | +++ | +++ | +++ | +++ | +++ | +++ | +++ | +++ | +++ |
| M4317 | +++ | +++ | +++ | +++ | +++ | +++ | +++ | +++ | +++ |
| M26796 | +++ | +++ | +++ | +++ | +++ | +++ | +++ | +++ | +++ |
| M4921 | +++ | +++ | + | + | +++ | +++ | - | +++ | - |
| M24531 | +++ | +++ | +++ | +++ | +++ | +++ | - | + | - |

## Discussion

We analyzed 2,088 lncRNAs in leaves of *B. rapa*; their characteristics such as number of exons, length of transcripts (except for NATs), and lower expression levels were similar to those reported in other plant species [40–45]. The total number of lncRNAs seems to be similar among species [46,47], but there tends to be low sequence conservation between different species [48]. Lower conservation of lncRNAs than mRNAs was observed between *B. napus* and its two ancestral species, *B. rapa* and *B. oleracea* [29,42], even though the hybridization was relatively recent event, ∼1,910–7,180 years ago [49]. We also found a lower sequence conservation of *B. rapa* lncRNAs among other species of the genus Brassica (*B. nigra, B. oleracea, B. juncea*, and *B. napus*). In 12 selected highly conserved lncRNAs, we found conservation of the lncRNAs at both the sequence and transcriptional level between *B. rapa* and *B. oleracea*, but there was transcriptional variation within species agreeing with the IRR diversification of each species after the allotetraploidization of *B. napus* [50]. The analysis of sequence homology of lncRNAs in *B. rapa* with other members of Brassica genus may deepen the understanding of the evolutionary dynamics of lncRNAs in the genus Brassica.

We found over 55% of each lncRNAs (65.0% lincRNAs, 55.0% NATs, and 71.7% incRNAs) overlapped with IRRs including TEs throughout the *B. rapa* genome. TEs are considered a source of siRNAs [51], and lincRNAs identified in *A. thaliana*, rice, and maize, were associated with 22.2%, 49.7%, and 51.5% of TEs, respectively [52]. In this study, we found 18.7% lincRNAs, 14.0% NATs, and 17.4% of incRNAs covered unique-mapped 24-nt siRNAs, and over 80% of lncRNAs covering the genomic regions encoding for 24-nt siRNAs overlapped with IRRs with perfect sequence match, suggesting that lncRNAs covering IRRs are also the likely source of siRNAs in *B. rapa*. For mRNA, the average expression levels of lncRNAs having 24-nt siRNAs was higher than that of lncRNAs not having 24-nt siRNAs [36]. LncRNAs and 24-nt siRNAs are known to be involved in the increase of DNA methylation, which agrees with reports of increased gene expression when DNA methylation overlaps with an exonic region [53]. The detailed mechanism is not clear, but our results also agrees that lncRNAs may be involved in this gene regulation mechanism.

The epigenetic functions of different lncRNAs have been identified in diverse organisms [21,22], but there is limited information in *B. rapa*. LncRNAs show a close association with DNA methylation of the genomic region encoding the lncRNA [54], but studies have focused more on the model plant species, *A. thaliana* [11,55]. In this study, we found a similar level of DNA methylation in regions encoding for lncRNA compared to previously described DNA methylation in the *B. rapa* genome [36]. Levels of DNA methylation can be positively regulated by siRNAs through the RdDM pathway [56], and Pol IV is important for biogenesis of 24-nt siRNAs [57]. Identification of long Pol IV-dependent transcripts is difficult because these transcripts can be rapidly processed to produce 24-nt siRNAs. There are several reports that detected Pol IV-dependent siRNA-precursor transcripts in *A. thaliana* of different lengths [57–59]. Most Pol IV-dependent transcript regions overlapped with Pol IV-dependent siRNA loci, and CHH methylation depends on the production of Pol IV-dependent transcripts [59]. However, if Pol IV-dependent transcripts are short as 30-40 nucleotides [57,58], the lncRNAs that we identified are of much longer length and not likely to be Pol-IV dependent transcripts that act as siRNA-precursors. Pol V transcripts accumulate at very low levels and have been difficult to identify. However, using RNA immunoprecipitation, Pol V-dependent lncRNAs were identified, and CHH methylation and 24-nt siRNA accumulation were shown to be restricted to Pol V transcribed regions [60]. In this study, about 13 % of lncRNAs overlapped with genomic regions encoding for 24-nt siRNAs, and the DNA methylation level of those regions was higher than the average. It has been reported that the median length of Pol V transcripts was 689 nt [60]. The length of the lncRNAs overlapping genomic regions encoding for 24-nt siRNAs identified in this study resembles the length of the Pol V transcript precursors.

The level of H3K27me3 plays a role in tissue specific gene expression [61]. LncRNAs are also considered to be involved in regulating histone modifications. Binding of lncRNAs to PRC2 has been observed in human/animal [62,63], and COLDAIR and COLDWRAP have been shown to recruit the PRC2 complex to *FLC* during vernalization in *A. thaliana* [13,14,64,65]. EMF2B is a component of the PRC2 complex and in rice, a mutant of *emf2b* was reported to lose H3K27me3 and derepress some of the lincRNAs, suggesting that expression of these lincRNAs is regulated by PRC2-mediated histone methylation of the region encoding the lncRNA [66]. We found that about 10-16% of lncRNAs overlapped with H3K27me3 regions, which is lower than the overlap with mRNAs. The pattern of H3K27me3 in lncRNA overlapping regions was similar to mRNAs; H3K27me3 accumulated in the body region of transcripts, especially around the transcription start site. However, there was no negative relationship between the presence of H3K27me3 and lncRNA expression levels, and incRNA coding regions having H3K27me3 marks showed higher expression levels than incRNA coding regions without H3K27me3 marks. H3K27me3 over lncRNA coding regions was also found in different tissues of maize, which was responsible for the regulation of tissue-specific lncRNAs expression [67]. Coincidentally, DNA methylation in intronic regions is known to influence splicing patterns [68], which may reflect epigenetic tissue specific gene regulation involving RdDM and H3K27me3. However, as our study focused on the analyses in a single tissue/developmental stage, we could not identify the lncRNAs that are transcriptionally silenced by H3K27me3, leading to the underestimation of lncRNAs with H3K27me3 marks. Further study will be required to examine the transcriptional regulation of lncRNAs by H3K27me3.

This study revealed that a small proportion of lncRNAs in *B. rapa* are conserved with other Brassica species. The majority of lncRNAs in *B. rapa* overlap with IRRs and there is some overlap with DNA methylation and 24-nt siRNAs, hinting at regulation of lncRNA expression through the RdDM pathway. Interestingly, some of the lncRNAs that overlapped with the genomic regions having H3K27me3 marks were more highly expressed, which was unexpected because of the known gene suppression activity of H3K27me3. This may indicate an unknown regulatory mechanism of lncRNAs by H3K27me3. However, the current study focuses on the genome-wide quantification of lncRNAs, 24-nt siRNAs, DNA methylation, and H3K27me3, and further quantitative analysis at the locus specific level may reveal a clearer relationship between these marks. Epigenetic regulation of the genome including lncRNA, 24-nt siRNAs, DNA methylation and H3K27me3 depend on the stage of development, and abiotic stress conditions. Further exploration of lncRNAs under different tissues and conditions may reveal lncRNAs with specific functions.

## Acknowledgement

We are grateful to Ms. Tomoko Kusumi for her technical assistance throughout this project.

## Supporting information

**S1 Fig.**
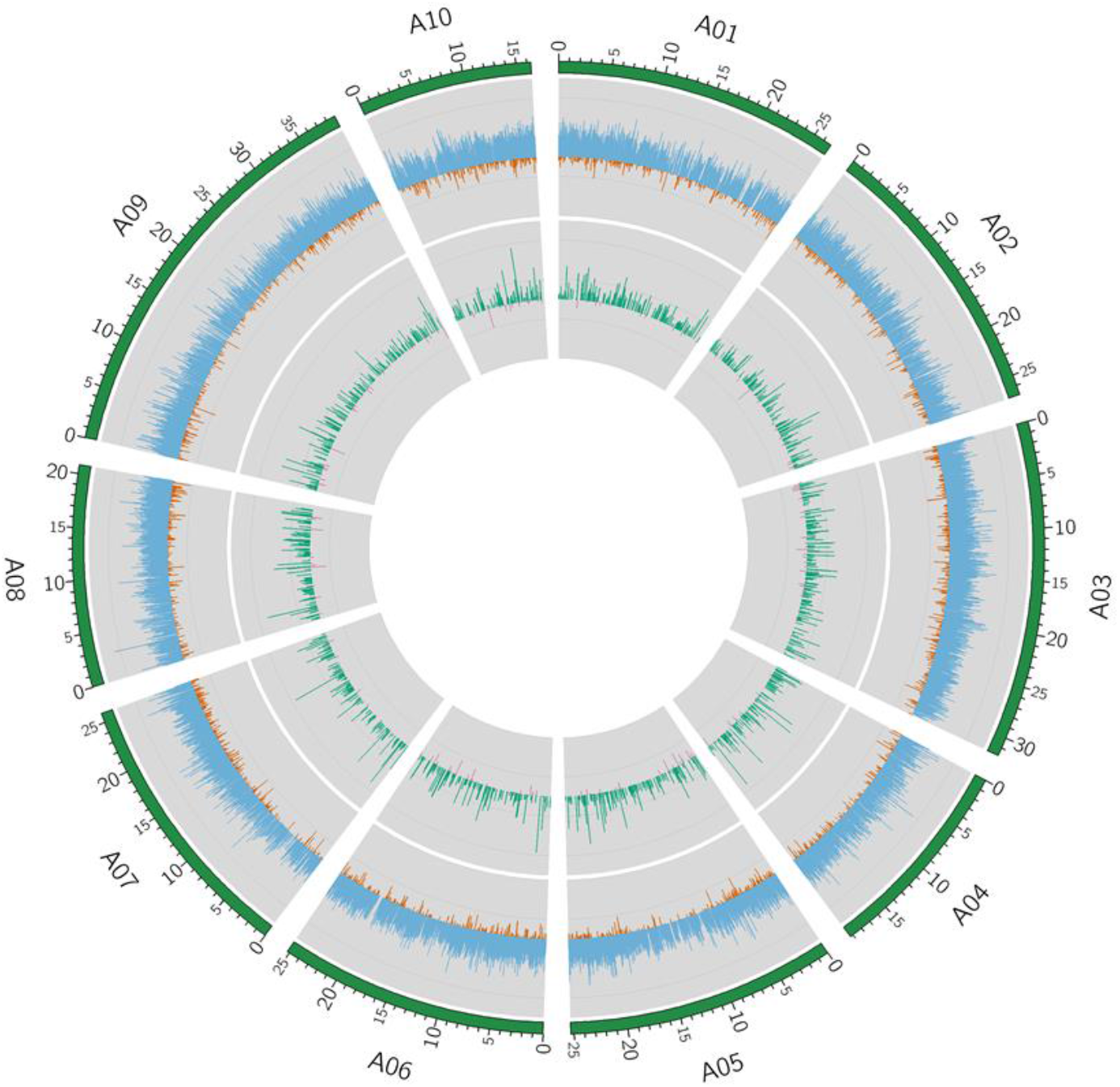
Chromosomal (A01-A10) distribution of lncRNA and mRNAs. Log_2_ FPKM of mRNAS represented in blue (positive values) and orange (negative values), and lncRNAs in green (positive values) and pink (negative values).

**S2 Fig.**
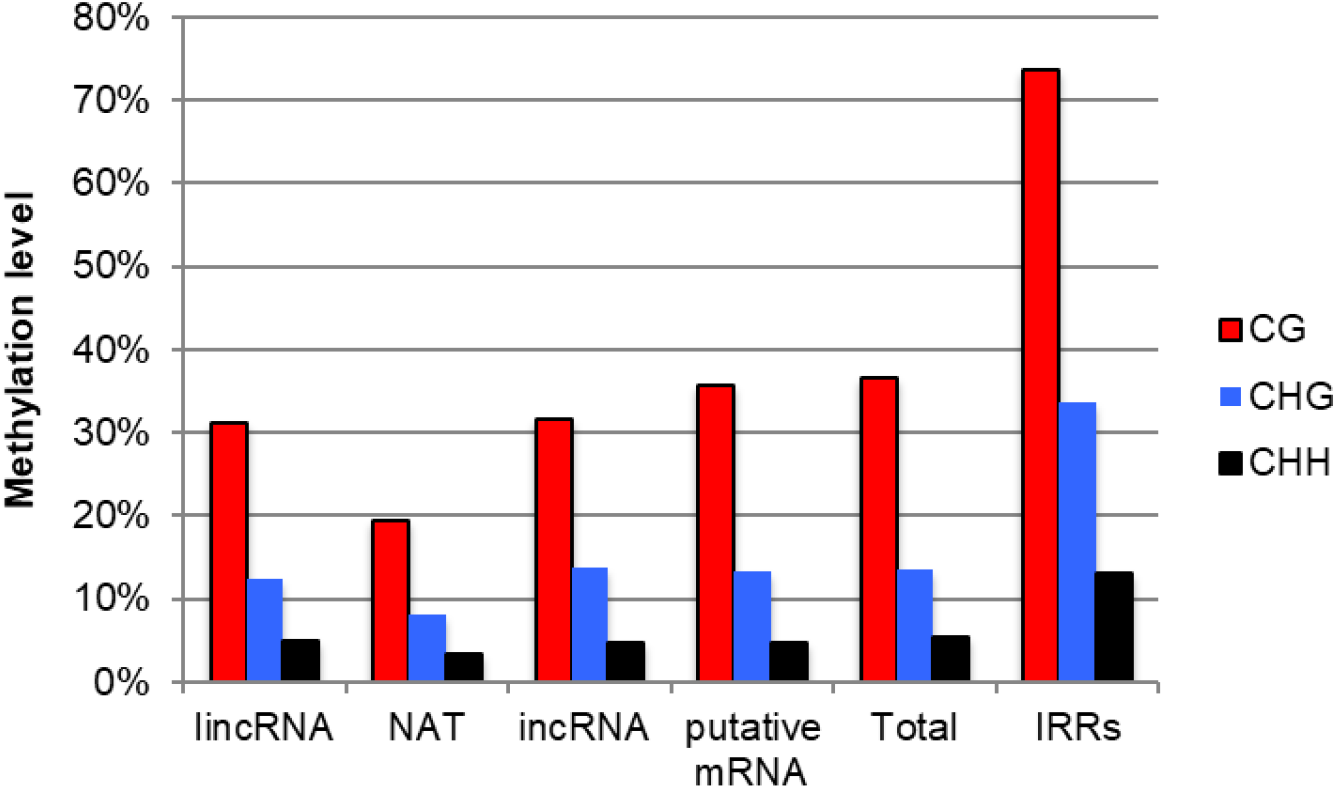
DNA methylation level of each type of lncRNAs, putative mRNA, total, and IRRs. “Total” represents the methylation level of all regions.

**S3 Fig.**
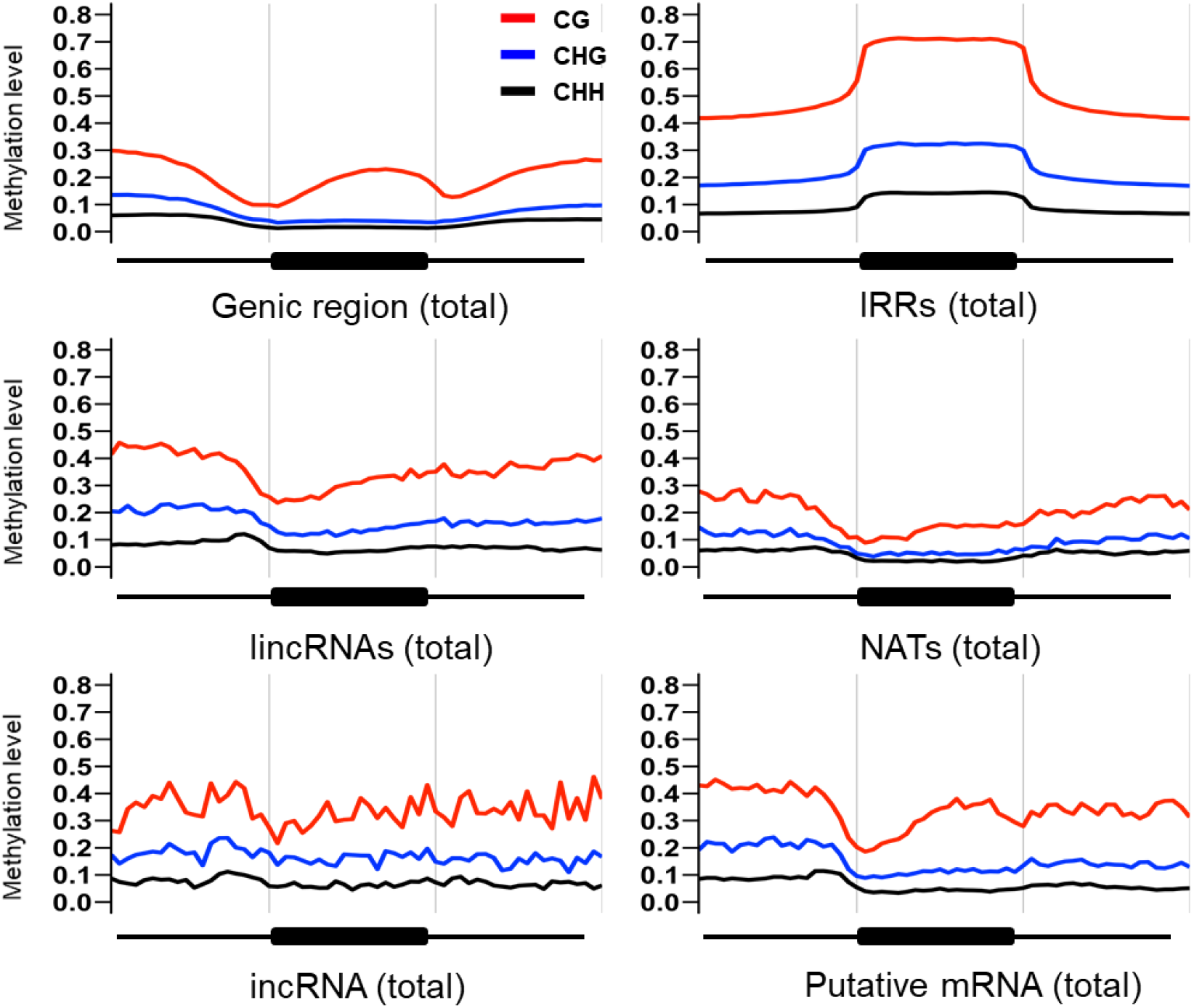
DNA methylation of the genic region, IRRs, or regions coding for putative mRNA and lncRNAs.

**S4 Fig.**
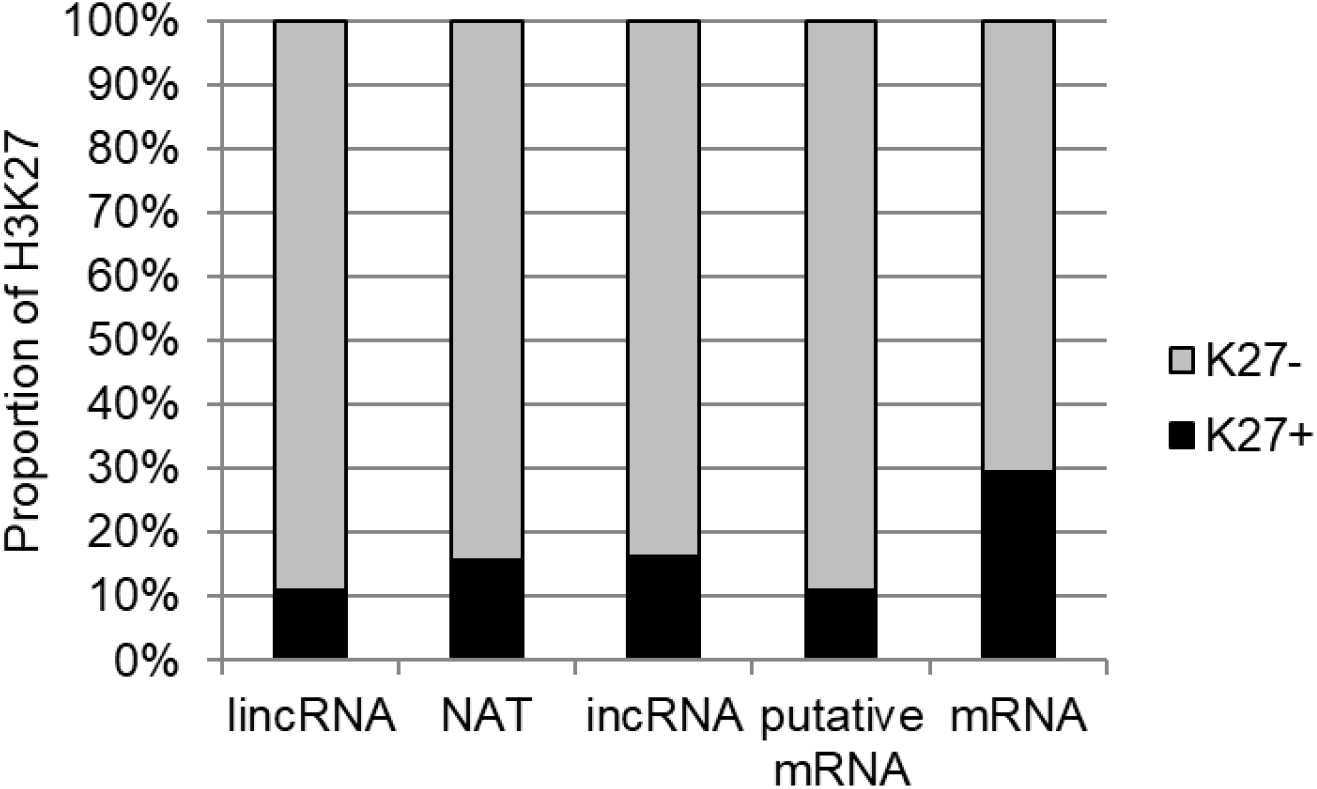
Proportion of lncRNAs, putative mRNA, mRNA with (K27+) or without (K27-) H3K27me3 on encoding regions.

**S5 Fig.**
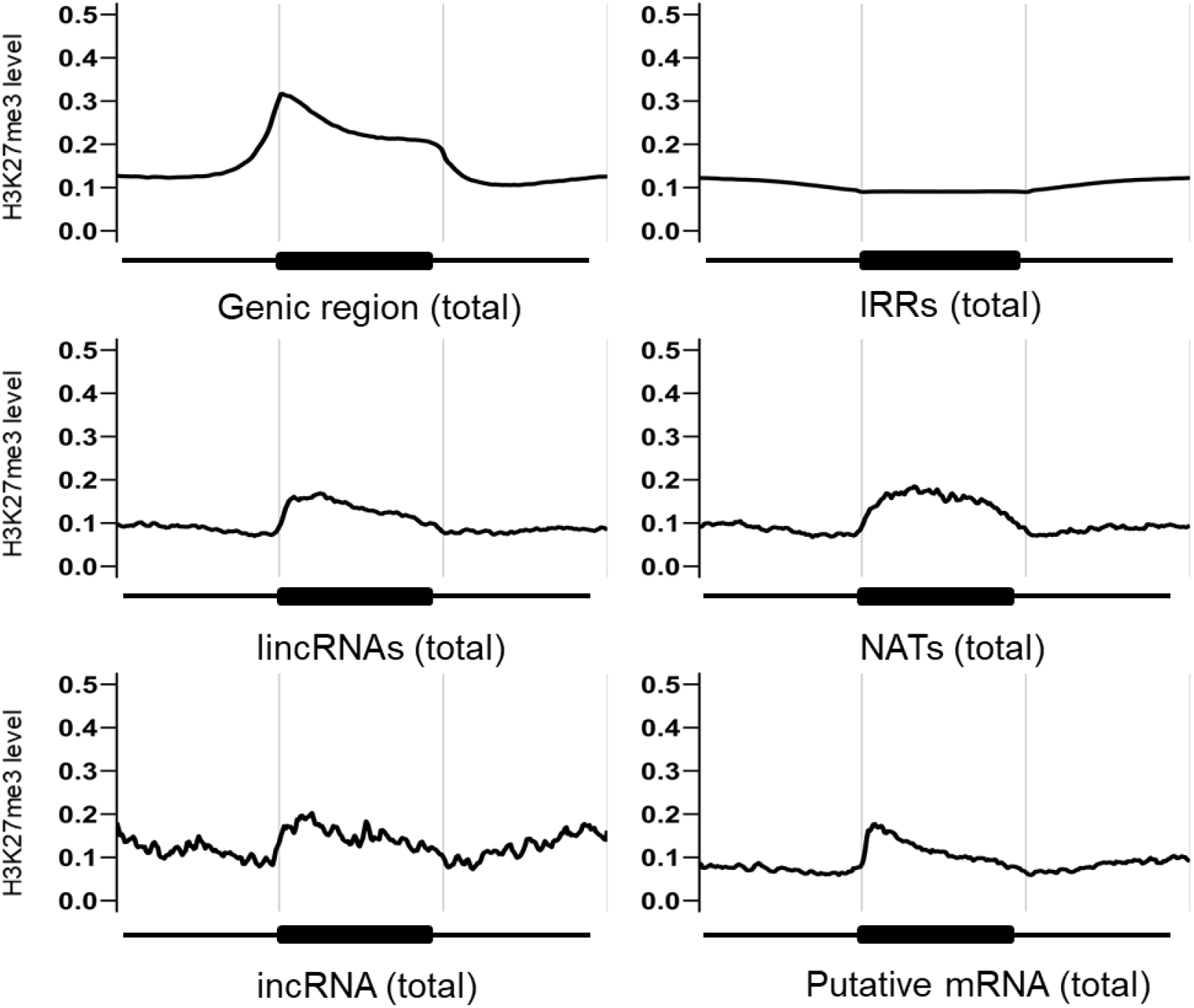
H3K27me3 mapped to the genic region, IRRs, or encoding regions of putative mRNA or lncRNAs.

